# Identifying interactions in omics data for clinical biomarker discovery using symbolic regression

**DOI:** 10.1101/2022.01.14.475226

**Authors:** Niels Johan Christensen, Samuel Demharter, Meera Machado, Lykke Pedersen, Marco Salvatore, Valdemar Stentoft-Hansen, Miquel Triana Iglesias

## Abstract

The identification of predictive biomarker signatures from omics data for clinical applications is an active area of research. Recent developments in assay technologies and machine learning (ML) methods have led to significant improvements in predictive performance. However, most high-performing ML methods suffer from complex architectures and lack interpretability. Here, we present the application of a novel symbolic-regression-based algorithm, the QLattice, on a selection of clinical omics data sets. This approach generates parsimonious high-performing models that can both predict disease outcomes and reveal putative disease mechanisms. Due to their high performance, simplicity and explicit functional form, these biomarker signatures can be readily explained, thereby making them attractive tools for high-stakes applications in primary care, clinical decision making and patient stratification.

## Introduction

### Background

The rapid increase in biological data obtained through high-throughput technologies offers new opportunities to unravel the networks of molecular interactions that underlie health and disease [1]. An important contribution to this is made by genomics, transcriptomics, proteomics, lipidomics and metabolomics studies, which generate thousands of measurements per sample and offer the unique opportunity to uncover molecular signatures associated with a particular condition or phenotype. These signatures have the potential to act as biomarkers, i.e. a biological characteristic used in the evaluation of normal, abnormal or pathogenic conditions. Biomarker profiles have been found to be particularly useful for medical decision making, where use cases such as surrogate endpoints, exposure, diagnosis and disease management have been identified [2]. Although the large amount of omics data contains extensive information, it is not always trivial to extract actionable insights from it. Challenges include the high dimensionality of datasets where the number of variables far exceeds the number of samples, unbalanced measured outcomes (target variables), heterogeneous molecular profiles with multiple subtypes of patients and diseases, and instrumental and experimental biases [3–5].

Classical statistical modelling has long been the gold standard for the analysis of genomics and transcriptomics data analysis. As a result, a significant amount of post-processing is required to condense information into meaningful results, e.g. through manual searching, enrichment or pathway analysis. An inherent challenge in the wide data matrices typical of omics is the existence of dependencies between features. This phenomenon is called “multicollinearity” or “concurvity” when linear and nonlinear dependencies are involved, respectively [6]. The increasing availability of affordable computing power and high-throughput omics data has led to the increasing use of machine learning (ML) in the life sciences and pharmaceutical industries. In addition, ML methods have been used for biomarker discovery based on omics data, where they are beginning to outperform state-of-the-art assays [7].

Due to the inherent noise of biological data and the “curse of dimensionality” [8, 9] (more features than observations), it is a non-trivial task to perform traditional machine learning without misleading or overfitting the model during training, such that it is unable to robustly predict outcomes on unseen samples [10]. In addition, most state-of-the-art machine learning models are difficult to interpret and are therefore often considered complex “black boxes” [11]. Applying black-box machine learning models such as random forests and neural networks to omics data has proven effective in identifying predictive biomarkers [12], but the underlying relationships between features remain hidden, and especially for decisions where the stakes are high, it has been argued that interpretable methods should be used wherever possible [13].

### Symbolic regression and parsimonious models

Recently, the QLattice, a new machine learning method based on symbolic regression (SR), has shown promising results in terms of performance and interpretability [14]. The goal of any implementation of symbolic regression is to model a relationship between one or more independent variables *X* and a dependent variable *y* by finding a suitable combination of mathematical operators and parameters. Even when considering only expressions with finite length, the search space is usually too large for any kind of brute force strategy, and thus, alternative methods are required. All SR algorithms can be thought of as methods of searching this combinatorial space effectively. Symbolic regression is an active field of research and there are multiple examples of recent implementations [15–18].

Symbolic regression is particularly suitable for scenarios where the number of features in the model should be small and their interpretation and interactions are of primary interest. Furthermore, it seeks to solve problems where the mathematical form of the data generating process cannot be assumed, or approximated, *a priori*. This is in contrast to the typical regression problem where parameters are fitted to a presupposed model, like linear models or polynomials. Thanks to its unconstrained nature, SR can usually attain higher performances while keeping the number of explicit parameters as low as possible.

It is well known that most functions can be approximated by using an arbitrarily large number of coefficients and functions belonging to a complete set (e.g. Fourier series, Chebyshev polynomials etc.). Analogously, one can theoretically build a model that explains *y* in terms of *X* with arbitrarily low train error, even if the approximated mathematical model is ostensibly different from the data-generating process. This does not necessarily pose a problem to types of research where the primary objective is to produce a working model that fits well the data, but vital information may be lost along with interpretability as model complexity grows. The most well-known example of this is deep neural networks, where e.g modern language models contain billions of parameters [19], inevitably trading off interpretability for performance.

In contrast, the aspiration of SR is that domain knowledge can be applied and extracted more efficiently by seeking simpler mathematical models to preserve explainability from a human perspective. In principle, this increases the likelihood of discovering driving mechanisms in data, and inclines SR methods toward maximum information gathering, which is vital in (e.g. life) sciences where both performance and interpretability is important. In practice, SR methods achieve this by using parsimonious models that explain the data with a minimal number of parameters. Additionally, one can use complexity measures such as Bayesian information criterion (BIC) and Akaike information criterion (AIC) to ensure that the resulting models generalize well from train to test set.

Here, we applied the QLattice to four different omics problems to identify biomarker signatures that predict clinical outcomes while also revealing new interactions in the data. We demonstrate how highly complex problems can quickly be condensed into a set of simple models that can be reasoned and used as hypotheses for potential mechanisms underlying the problem at hand.

## Methods

### The QLattice

The QLattice is a symbolic regression engine that aims to solve the optimization problem of finding the functions that best fit the data. It applies an evolutionary algorithm framework to find the combinations of inputs, operators and parameters that minimise the fitting error in a supervised learning problem (see [20] for seminal work on genetic programming, and a practical guide in [21]).

The QLattice algorithm works as follows: first it generates an initial sample of functions, fits them with gradient descent, and evaluates them for fitness. This initial sample is formed using a set of estimated priors assigned to each input based on its mutual information with the output. Then, the best performing functions are used to create a new generation of functions consisting of three groups: 1) the best performers from the previous generation 2) mutated versions of the best performers from the previous generation and 3) a completely new set of sampled functions. Instead of sampling mutations and new functions from a uniform distribution of inputs and operators, the QLattice draws from a probability distribution that is learned thanks to a mapping between the functional space and a lattice. Thus, with each generation the QLattice improves the probability distribution estimation. As this iterative process continues, the QLattice expands the search for the best fitting functions. The result of a training run is a list of functions sorted by a user-defined quality metric. These functions serve as hypotheses that each serve as their own solution to the problem. A more extensive description of the methodology can be found in [22].

The QLattice can be used in both regression and classification tasks for supervised learning problems. In the case of classification, the algorithm is designed to work with binary problems, although it can be easily extended to multi-class targets using a one-versus-rest approach (see [23] for detailed description of of the method). All the QLattice models discussed in this manuscript are trained to perform binary classification tasks. The target variables are encoded as 0 or 1, and the output of the models is to be interpreted as a probability. In order to keep the outputs between 0 and 1, all the mathematical expressions are wrapped with the logistic regression function 1/(1 + exp(−*f* (*X*))), expressed throughout the text as *logreg*.

The Feyn Python library [24] is the interface between the user and the QLattice, and it is used to train and analyse new models. Its high-level train function returns a list of ten models sorted by a criterion of choice (see documentation [25]). The default sorting option is the Bayesian information criterion (BIC), which amounts to the training loss plus a complexity penalty, and allows selection of the most generalisable models without compromising training speed [9].

A majority of the plots in this paper were created using the Feyn [24] (which uses Matplotlib [26] extensively), and the Seaborn [27] libraries.

### Cross-validation

Overfitting and spurious correlations are major concerns when applying machine learning to the wide datasets typical of many areas of computational biology such as genomics, transcriptomics, and proteomics (that is, when the number of features is much larger than the number of observations). For these kinds of datasets, simple models with complexity penalties tend to offer competitive performances [9]. This is the case of the models selected by the QLattice when the BIC criterion is enabled. The BIC criterion used for model selection, however, does not provide an unbiased estimate of the test performance.

Therefore, we use a standard k-fold cross-validation scheme to estimate the performance of the QLattice and determine what one can expect from the models selected by it. We use a scheme with 5 folds: 4 folds as a train set, and 1 as a test set. In each of the 5 training loops, we reset the QLattice and call the train function to avoid “data leakage” in the feature selection. Individual models’ performances are estimated using single train/test splits.

Finally, in the supplementary material section, we present a comprehensive benchmark of the QLattice along with other machine-learning algorithms in combination with different feature selection techniques on all four data sets discussed in the manuscript.

### Selection of models for further analysis

In machine learning the emphasis is usually put on test-set performance. In most instances, model selection is done with the sole goal of finding the models that will generalize best on new data. BIC is a good example of such a model selection tool, and the QLattice uses it to explore and find the models with strongest signal – both in the train and test sets. In the cases where the user is only interested in prediction performance one should select the model with the lowest BIC score. This is the selection criterion we followed in the benchmarking section.

Having said that, interpretable algorithms like the QLattice have more goals than performance. They are also used to generate hypotheses about the features involved in a process, and their specific relations. When using BIC as a criterion, all the models returned by the QLattice can be expected to highlight robust patterns in the data. Although the performance of the models might differ, one should consider all of them valid. The list of models returned by the QLattice might highlight a combination of patterns: different mechanisms, the same one represented by multicollinear features, already known biomarkers, or completely new candidates. The evaluation of the user (*human in the loop*) is then necessary to extract the relevant learnings from the models, and put them in the context of the question at hand.

For the sake of clarity, we only discuss one model in each of the first three cases. We selected the models according to performance and interpretability: when models had very similar performances, we chose the simplest models first. In the insulin response case we used evidence from previous studies to choose a model, where the gene features could be easily interpreted. We conclude by noting that the biomarker candidates selected by the models on the different cases analysed in this paper should be further investigated – both in relation to the diseases’ mechanisms and whether they might come from confounding factors (e.g. cohort dependencies).

### Data preparation

#### Proteomics: Alzheimer’s disease

The data was taken from Bader et al. [28] and consists of 1166 protein expression of the cerebrospinal fluid of 137 subjects, collected in three sample groups (we address the possible confounding factors in the results section). We used the QLattice to predict whether a patient would develop Alzheimer (dependent variable = 1) or not (dependent variable = 0).

#### Genomics: Relevant genes for insulin response in obese and never-obese women

The data was retrieved from Mileti et al [29]. The dataset consists of gene expression from a total of 23 never obese and 23 obese women sequenced before and 2 years after bariatric surgery (post obese) using RNA sequencing (CAGE) [29]. The only pre-processing done was to normalise the data from raw counts to TPM (tag-per-million normalisation, the gold standard for CAGE data [29]). The data is balanced for MValue, BMI, and Age across sample groups. We used to QLattice to model the response to insulin based on gene-expression measurements and predicted whether an individual is in a fasting (target variable = 0) or hyperinsulinemic (target variable = 1) state.

#### Epigenomics: Hepatocellular carcinoma

The data was processed to contain only the 1712 most important features, filtered for variance. The curated dataset contained 1712 CpG island (CGI) features with a binary target of 55 cancer-free (target variable = 0) and 36 cancer (target variable = 1) individuals coming from a single sample group (plasma samples) [30]. The CGI features cover the methylated alleles per million mapped reads.

#### Multi-omics: Breast cancer

The data set was obtained from Ciriello et al. [31] and contains multi-omics data from 705 breast tumor samples of different patients. We use the QLattice to predict outcomes; a survival outcome is encoded with target variable = 0 (611 patients) and a fatal outcome with target variable = 1 (94 patients). The data was extracted from The Cancer Genome Atlas through the R-package “curatedTCGAData” [32] and included 4 data types: somatic mutations, copy number variations, gene expressions and protein expressions. The raw data was pre-processed with a variance threshold limiting each type of input to the highest variance features. The data was stratified for lobular and ductal subtypes in each train/test split and the models were assessed for potential confounding influences from factors such as age, stage and treatment regimen.

## Results and discussion

In the following cases we showcase different aspects of the QLattice using 4 different omics data types:

- **Interpretability:** In the proteomics case, we show how the QLattice finds high-performing models that can be easily interpreted.
- **Feature combinations:** In the genomics case, we demonstrate how the QLattice finds biomarker signatures that together explain the data better than any single feature on its own.
- **Multicollinearity:** In the epigenomics case, we show how the QLattice deals with multicollinearity typical of omics data by chosing the combination of features that best explains the target while minimising complexity of the model.
- **Non-linear interactions:** In the multiomics case, we highlight how the QLattice can find non-linear interactions within and across omics dataypes that help to stratify patient populations.

### Proteomics: Alzheimer’s disease

#### Background

Despite many decades of research, neurodegenerative diseases remain a major threat to human health and are a substantial cause of mortality. Alzheimer’s disease (AD) is the most common type of dementia, and currently no therapeutics can halt or significantly slow its fatal progression [33]. Furthermore, short of an autopsy, there is no definitive way to diagnose AD and it is in general impossible to predict who will develop the disease.

Here, we demonstrate how the QLattice can be used to discover protein biomarkers for AD working with the data from [28].

We will use this example as an introduction to the QLattice functionality and capabilities.

#### Model analysis

After splitting the dataset into 80% train and 20% test partitions, we ran the QLattice on the train partition to obtain ten best unique models from the QLattice (Table 1). Each model points to a relation that serves as a data-derived hypothesis. Thus, all ten models potentially hold insights into the mechanisms involved in AD.

**Table 1.** The lowest BIC-scoring models returned by the QLattice for the AD dataset. The majority are linear and contain three features. Training set AUC performances are comparable.

| Functional form (logreg()) | N. Features | BIC | AUC Train |
| --- | --- | --- | --- |
| LILRA2 + MAPT + age at CSF collection | 3 | 46.11 | 0.98 |
| IGKV2D-29 + LILRA2 + MAPT | 3 | 49.11 | 0.98 |
| FAM174A + IGLV4-69 + MAPT | 3 | 49.28 | 0.97 |
| MAPT*(AJAP1 + SERPINE2.1) | 3 | 49.45 | 0.97 |
| ENOPH1 + GPC1 + MAPT | 3 | 51.42 | 0.97 |
| GPC1 + MAPT + age at CSF collection | 3 | 52.27 | 0.98 |
| ENDOD1 + MAPT + PPIA | 3 | 54.08 | 0.97 |
| GPC1 + MAPT + SERPINE2.1 | 3 | 54.86 | 0.97 |
| IGLV4-69 + MAPT | 2 | 54.95 | 0.97 |
| MAPT + NXPH3 + SPINT2 | 3 | 57.0 | 0.97 |

We chose the model with the lowest BIC-score for thorough analysis. This model uses MAPT, age at CSF collection, and LILRA2 as inputs combined with additions to predict the probability of AD for a given patient. As a machine learning model, it can be analysed in terms of test prediction metrics (Fig. 2) to verify that the relations found are not spurious (see [34] for a review on the matter).

We ran the cross validation scheme outlined in the Methods section. The estimated test performance of the QLattice top models was AUC = 0.94 (mean of the five folds, with a standard deviation of 0.05). We note that the predictive power might be over estimated due to the presence of confounders in the data.

#### Model Interpretation

The known AD biomarker MAPT (tau protein) was consistently found in the highest scoring QLattice models, while the additional features varied between models. Fig. 1 shows how MAPT contributes prominently to the signal of the chosen model. The plot shows the signal flow in the model, and the colour represents the strength of the association of each node to the clinical outcome. The association measure used is mutual information [35]. Thus, the features age at CSF collection and LILRA2 are both secondary to MAPT but both improve the model as made clear by the rising mutual information numbers displayed on top of the nodes. MAPT on its own has a mutual information score of 0.37 but this number rises to 0.56 when applying the additional features and the right mathematical operators – in this case additions.

**Figure 1.**
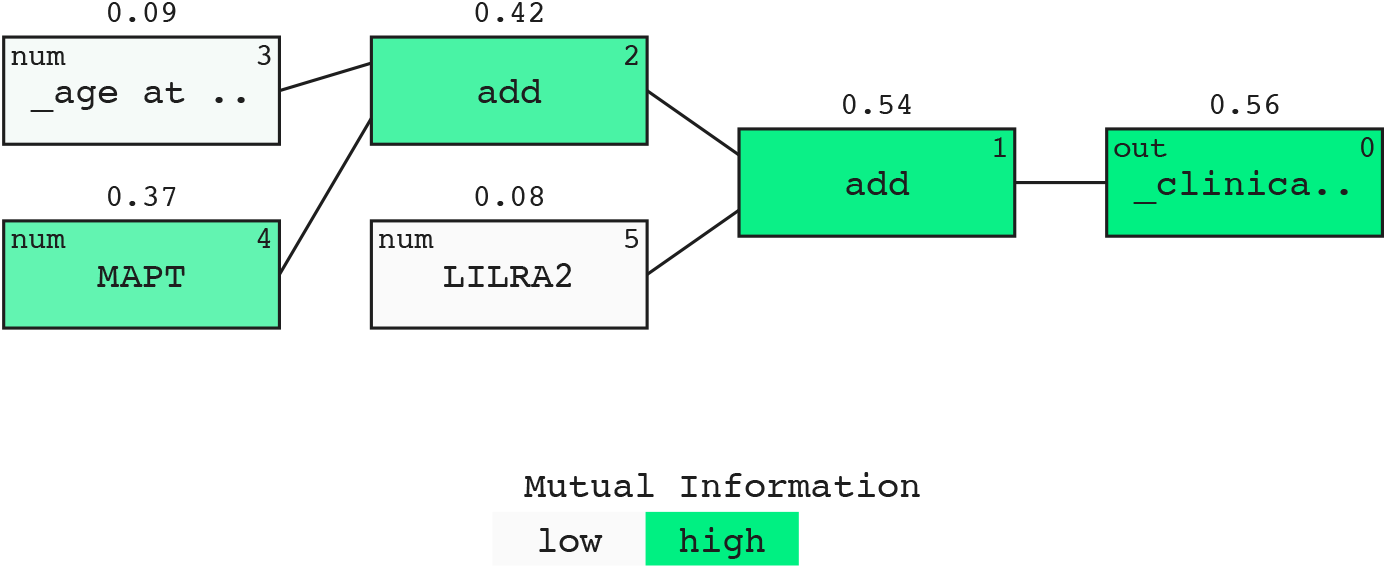
Model signal path for AD. A prominent signal contribution from MAPT was found in all 10 models (green). The signal is expressed in terms of mutual information and displayed above the nodes in the model (see [35])

The partial dependence plot (Fig. 3) shows that at fixed LILRA2, higher levels of MAPT leads to positive AD prediction. When the MAPT level reaches around 25,000 the model starts to predict AD-positive. In addition, the effect of age is displayed in the plot. Unsurprisingly, at a higher age comparably lower MAPT levels trigger the model to predict AD-positive (when the predicted probability rises above 0.5), as displayed by the coloured curves.

**Figure 2.**
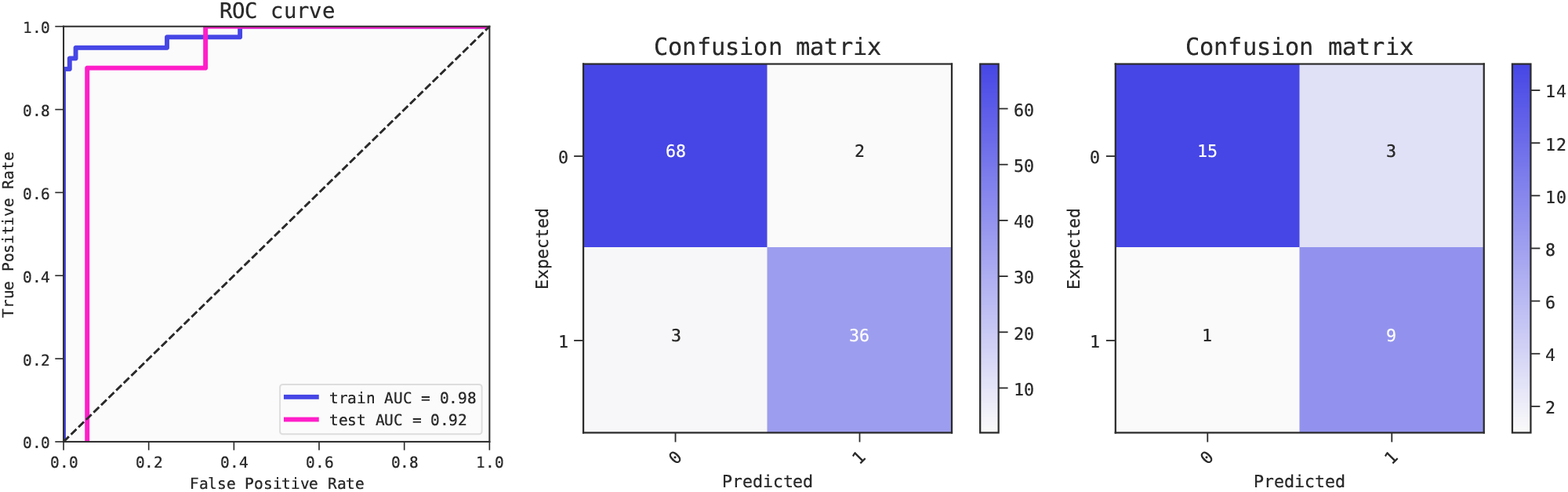
Metrics of the best model (ranked by BIC criterion) for predicting Alzheimer’s Disease. The model is robust as shown by the relatively small drop in performance from the training set (AUC 0.98) to the test set (AUC 0.92). Receiver operator characteristic (ROC) curves (left) and confusion matrices for training set (center) and test set (right).

**Figure 3.**
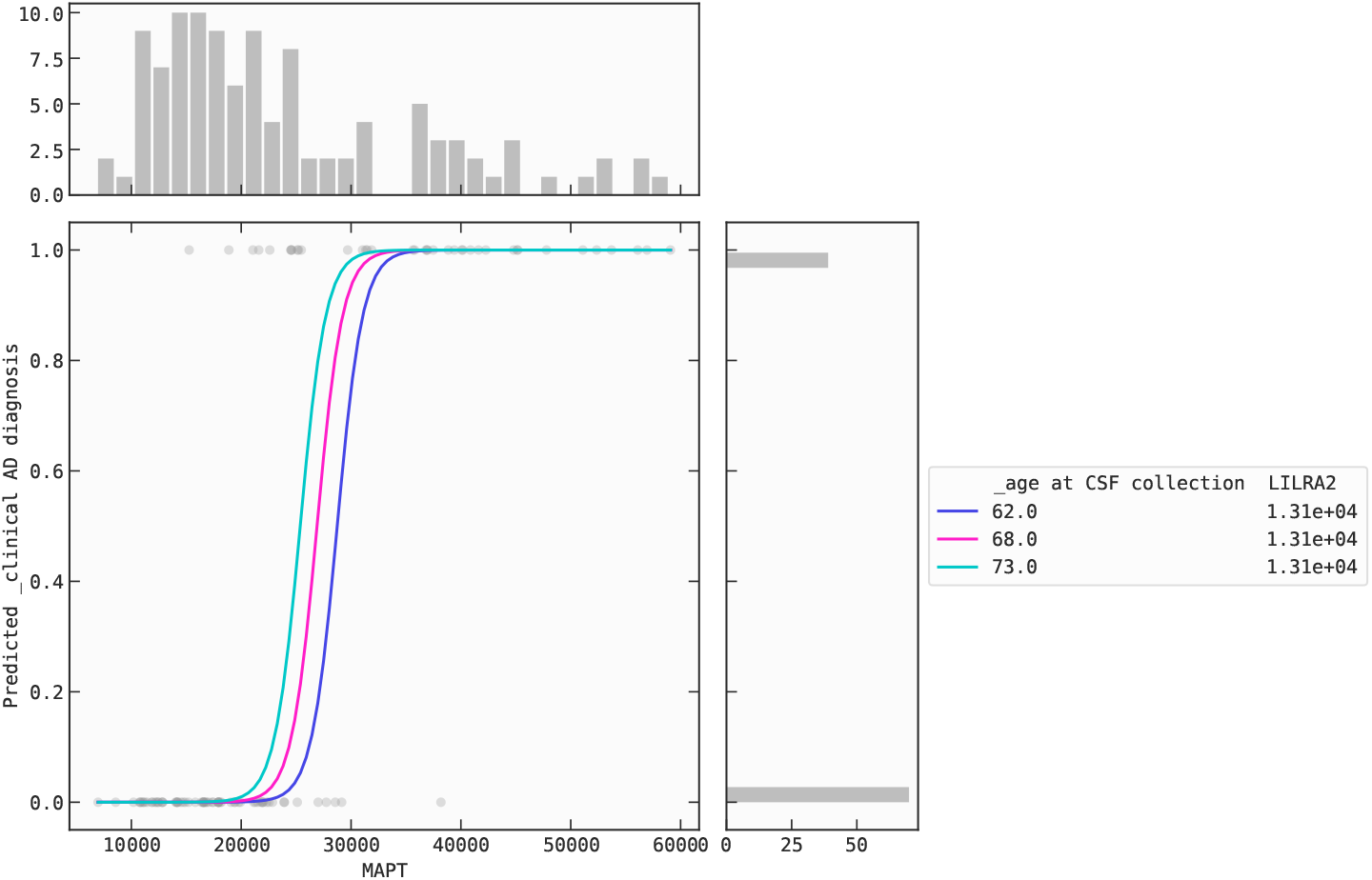
Partial dependence plot for the AD model: Marginal effect of MAPT on AD-risk.

It should be mentioned that the protein levels of LILRA2 only present a significant difference between AD and non-AD patients for one of the cohorts depicted in [28] (p-value of 0.008 with a Student’s t test). Since we fix LILRA2 in Fig. 3, the model response to MAPT is cohort-independent. Moreover, the three age values depicted in Fig. 3 are all between 62 and 72 years-old. This ensures that the observed model response is free from any bias arising from the younger control group present in one of the cohorts [28]. Including suspected confounding factors in the model is a standard practice to statistically control for confounders when using linear and logistic regression (see [36] for an extensive review on the topic).

The QLattice models provide data-derived hypotheses that can quickly provide an overview of possible explanations to a given question. For instance, the first model in Table 1 may be translated into the following hypothesis: “MAPT is a main driver of AD since it is positively correlated with AD status”, “MAPT interacts both with the age of the patient and with the protein LILRA2”. Thus, the mathematical simplicity of the QLattice models allows direct translation into hypotheses that can be readily understood and tested. This marks a significant departure from black-box ML models, where the inner working of the models is usually more opaque.

### Genomics: Relevant genes for insulin response in obese and never obese women

#### Background

Obesity is a major public health problem, and obese people are at higher risk of heart disease, stroke and type 2 diabetes. Obesity is considered a medical condition caused by eating more calories than necessary, but it can also be caused by a decreased response to insulin.

To shed light on this, a recent publication [29] focused on white adipose tissue (WAT), which is one of the main insulin-responsive tissues. In this study, obese subjects underwent gastric bypass surgery and lost weight. Weight loss can support the subsequent restoration of the insulin response. In [29], insulin sensitivity was determined using the hyperinsulinemic euglycemic clamp, while the insulin response was measured using cap analysis of gene expression (CAGE) from 23 obese women before and 2 years after bariatric surgery. To control for the effects of surgery, 23 never obese women were also included.

The experiment was designed to understand the effects of insulin on the expression of different genes. In traditional differential gene expression (DGE) analysis the individual genes with the strongest and most consistent changes between conditions are highlighted. Here, we propose a complementary approach to DGE analysis that uses the QLattice to identify sets of genes and their interactions that best separate two groups of samples.

Specifically, we modelled the response to insulin based on gene-expression measurements and predicted whether an individual is in a fasting or hyperinsulinemic state. As well as being a predictive algorithm, the QLattice looks for different interactions between genes that describe the insulin response in two classes of individuals.

#### Model analysis

We inspect the ten models returned by the QLattice in Table 2 after we ran it on the training set (80%-20% split). We select the second model for further analysis because it contains PDK4, an established insulin target [29]; C2CD2L, a positive regulator of insulin secretion during glucose stimulus; and PHF23 a negative regulator of autophagy. To our best knowledge, defects in autophagy homeostasis are also implicated in metabolic disorders such as obesity and insulin resistance as discussed in [37]. The high performance of this model is summarized in Fig. 4 for both the training and test sets. In addition, the QLattice identified other genes known to be insulin targets or found in the paper such as C19orf80 and LDLR [29].

**Table 2.**
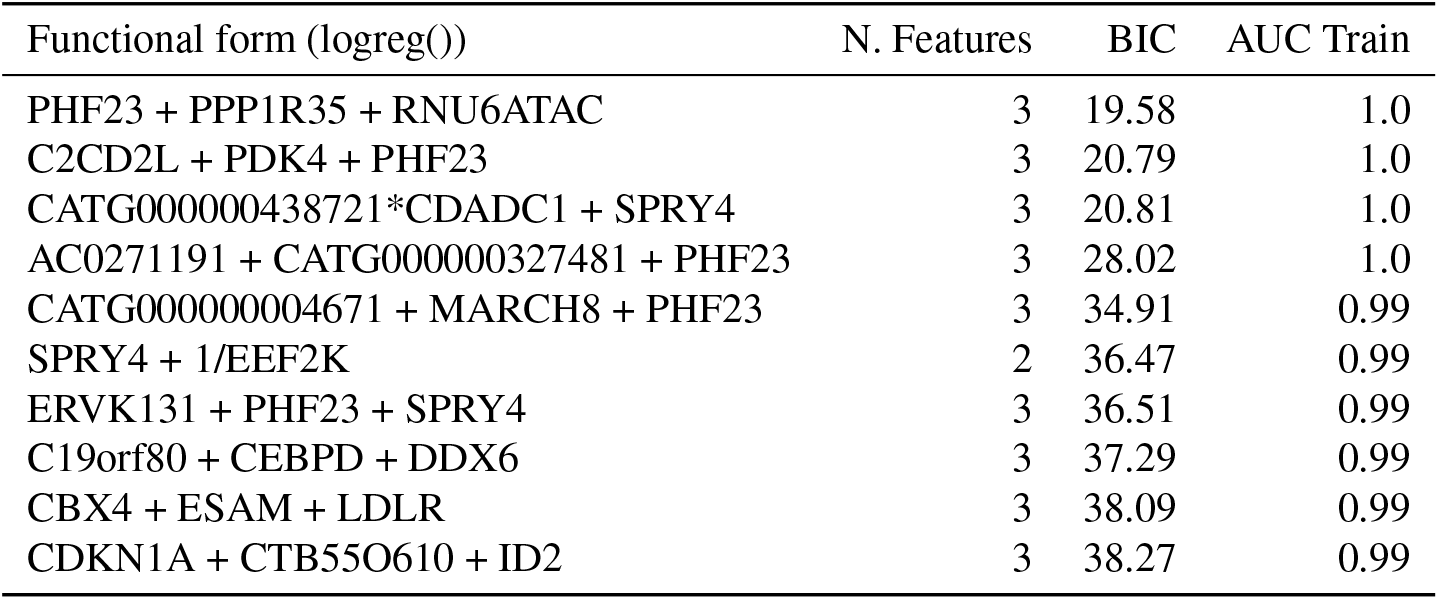
Lowest BIC-scoring models returned by The QLattice for the insulin response.

| Functional form (logreg()) | N. Features | BIC | AUC Train |
| --- | --- | --- | --- |
| PHF23 + PPP1R35 + RNU6ATAC | 3 | 19.58 | 1.0 |
| C2CD2L + PDK4 + PHF23 | 3 | 20.79 | 1.0 |
| CATG000000438721*CDADC1 + SPRY4 | 3 | 20.81 | 1.0 |
| AC0271191 + CATG000000327481 + PHF23 | 3 | 28.02 | 1.0 |
| CATG000000004671 + MARCH8 + PHF23 | 3 | 34.91 | 0.99 |
| SPRY4 + 1/EEF2K | 2 | 36.47 | 0.99 |
| ERVK131 + PHF23 + SPRY4 | 3 | 36.51 | 0.99 |
| C19orf80 + CEBPD + DDX6 | 3 | 37.29 | 0.99 |
| CBX4 + ESAM + LDLR | 3 | 38.09 | 0.99 |
| CDKN1A + CTB55O610 + ID2 | 3 | 38.27 | 0.99 |

**Figure 4.**
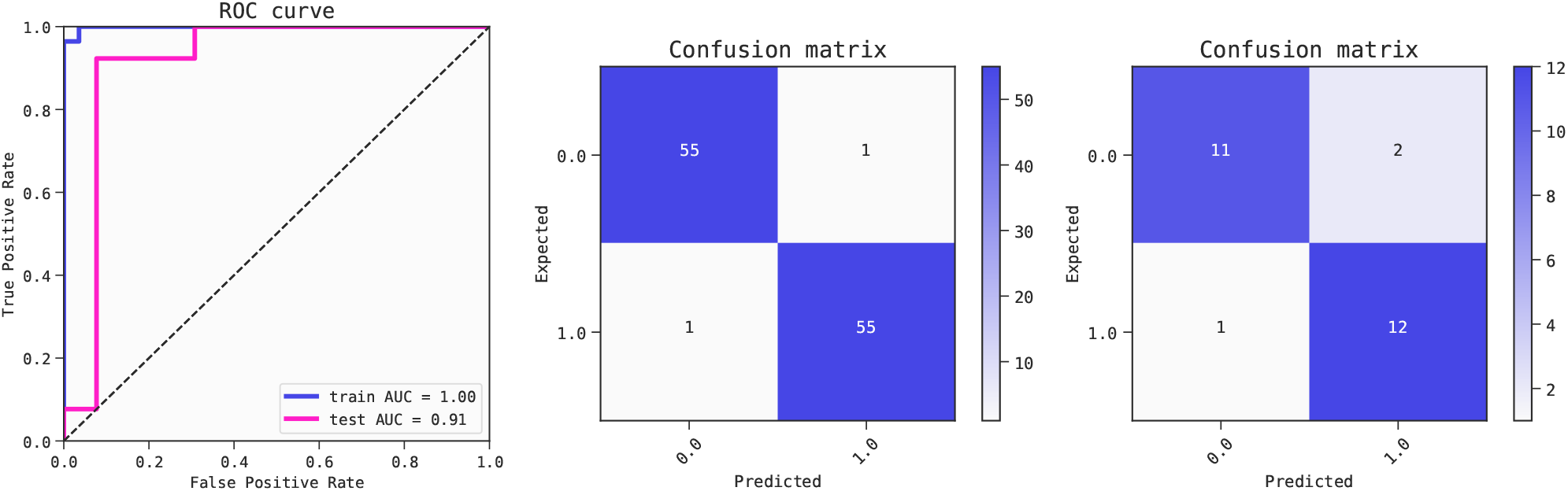
ROC AUC scores (left) for the selected three feature model for insulin response. Confusion matrices (center: train, right: test)

#### QLattice as complementary approach to differential gene expression

Differential gene expression analysis (DGE) is generally used to detect quantitative changes in expression levels between experimental groups based on normalised read-count data. There are several methods for differential expression analysis based on negative binomial distributions [38, 39] or based on a negative binomial model (Bayesian approaches) [40–42]. Differential expression tools can perform pairwise comparisons or multiple comparisons.

Alternatively, DGE can be used to identify candidate biomarkers, as it provides a robust method for selecting genes that offer the greatest biological insight into the processes influenced by the condition(s) under investigation. However, this robustness can sometimes translate into rigidity. Signatures expressed in linear combinations, interactions or through non-linear relationships may be overlooked when using DGE.

Symbolic regression based ML models offer a complementary view on the data and highlight predictive signatures. The advantage of this approach is that even simple feature combinations can lead to a high predictive performance. As we can see in the model decision boundaries of Fig. 5, a linear combination of the features PDK4, PHF23 and C2CD2L can characterize the insulin response for almost all individuals in the sample. The strength of the signal is found as well in the test set (see Fig. 4).

**Figure 5.**
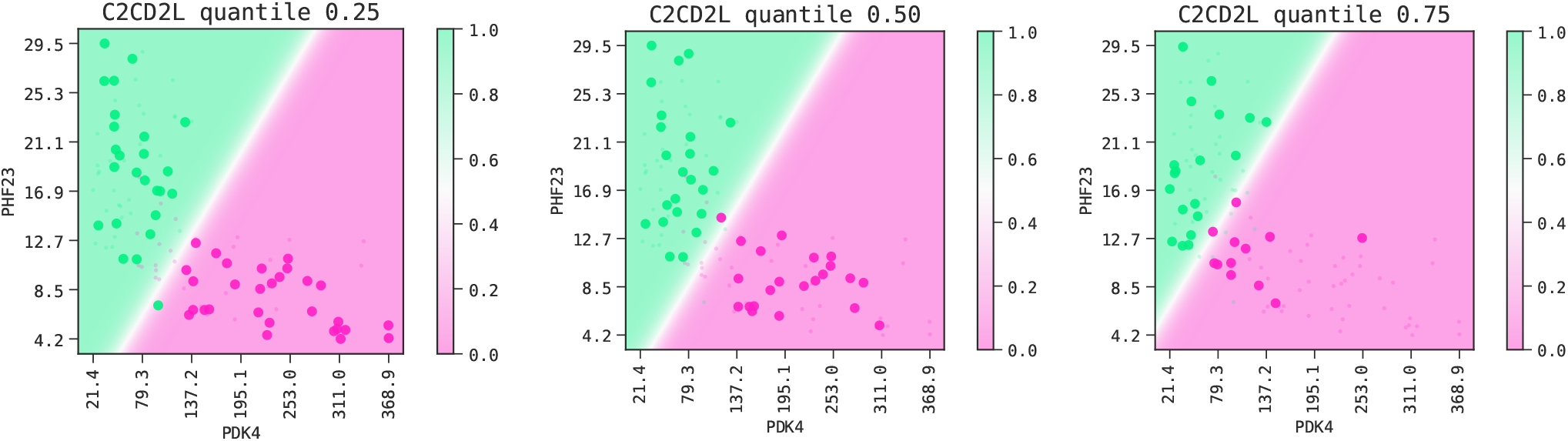
Decision boundaries of the selected model. We keep the feature C2CD2L fixed at the values corresponding to the 0.25, 0.50 and 0.75 quantiles.

Although PDK4 and PHF23 are reported as significant in the DGE analysis (according to FDR), they do not appear at the top of the list ordered by log-fold change (the one used in [29]). This apparent discrepancy between the DGE and the QLattice choice can be explained by the fact that the DGE only considers the univariate distributions. From the density plots in Fig. 6 we can indeed see a considerable overlap between the two classes when we look at the univariate distributions of PDK4 and PHF23, which is smaller for the distributions of the linear combination of genes. The effect can also be seen in the mutual information between the variables or their combinations, and the target variable.

**Figure 6.**
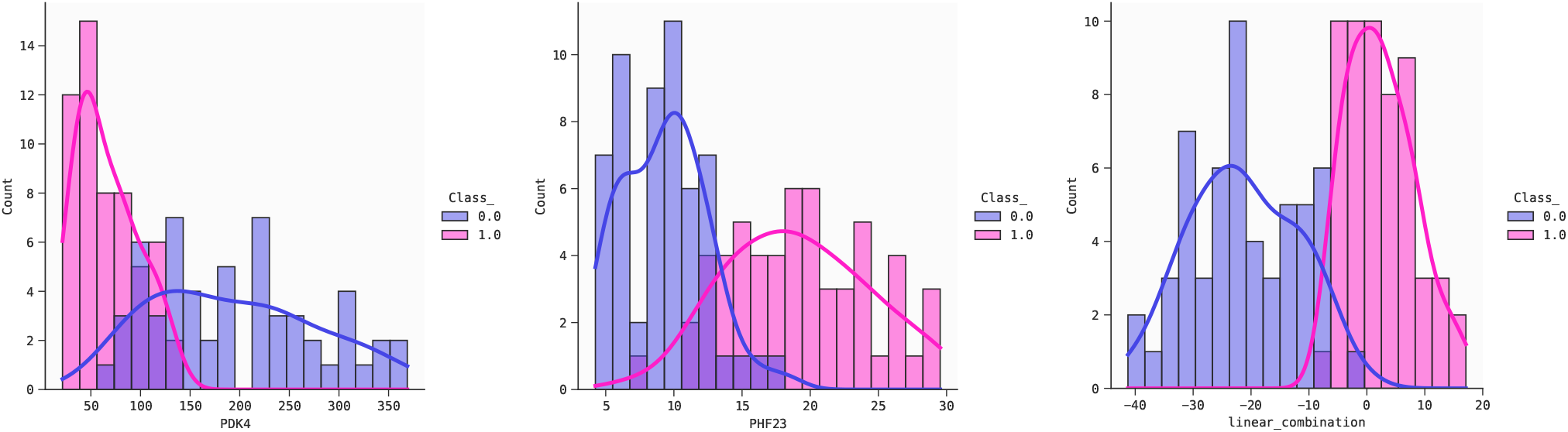
Distributions of the two classes for the variables PDK4 (left), PHF23 (center) and the linear combination found in the second model of Table 2 (right).

In summary, we find that the QLattice can be used as a complementary method to DGE, as it is very good at finding feature combinations that carry strong signals, and as it efficiently explores the feature space without requiring an exhaustive exploration of all features. There have been efforts in this direction using mutual information and partial information decomposition [43]. Consequently, the QLattice can suggest specific operations for the proposed combinations and help to better understand biologically relevant interactions that were previously hidden.

### Epigenomics: Hepatocellular carcinoma

#### Background

Primary liver cancer is a major health burden worldwide and develops in response to chronic inflammation of the liver. This can be caused by a number of insults such as viral infections as well as both alcoholic steatohepatitis (ASH) and non-alcoholic steatohepatitis (NASH). The most common form of liver cancer is hepatocellular carcinoma (HCC), which accounts for 90% of liver cancers and is the third leading cause of cancer mortality worldwide [44, 45].

In this HCC diagnosis example, we explore how the QLattice performs on highly multicollinear data. The dataset was taken from a study by Wen et al. [30] and contains data generated by methylated CpG tandems amplification and sequencing (MCTA-Seq), a method that can detect thousands of hypermethylated CpG islands (CGIs) simultaneously in circulating cell-free DNA (ccfDNA). The aim is to explain liver cancer occurrence using methylation biomarkers as features. After pre-processing (see Methods) the curated dataset contained nearly two thousand features. As demonstrated below, the QLattice gave highly predictive models using only a few key interactions.

#### Model analysis

As in the previous case, we split the dataset into train and test partitions (80%-20%) and ran the QLattice with default settings on the training set. We inspected the ten models returned in Table 3 balancing simplicity and performance. The ten models all perform equally well and we therefore chose the model with the least features for further examination (n. 5). Its performance metrics are summarized in Fig. 8.

**Table 3.**
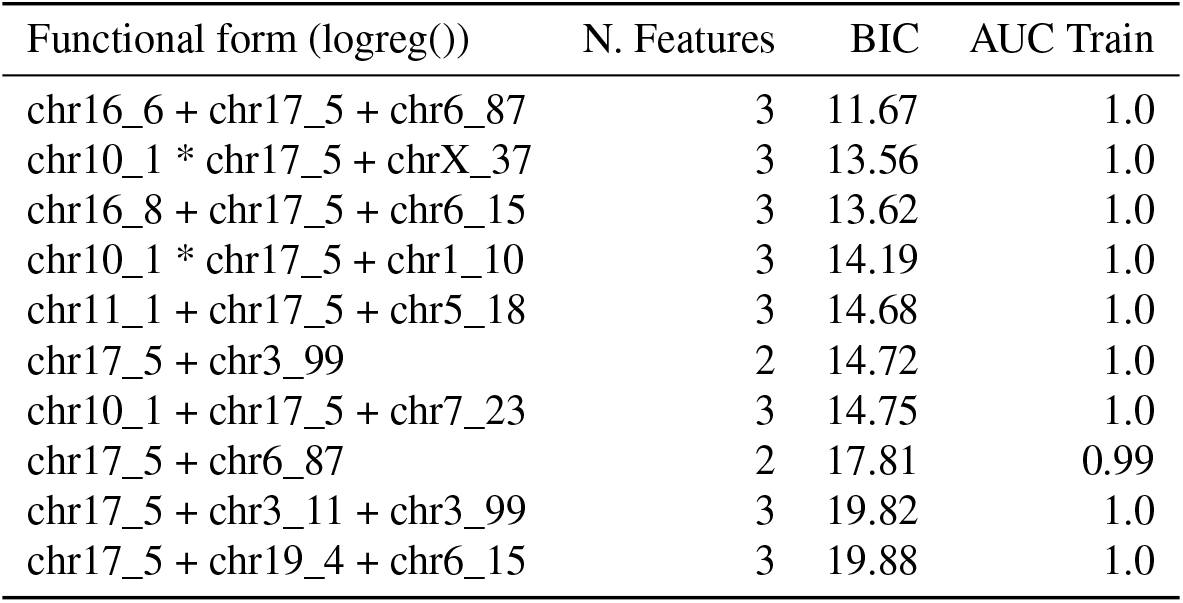
The lowest BIC-scoring models returned by the QLattice for the HCC dataset.

**Figure 7.**
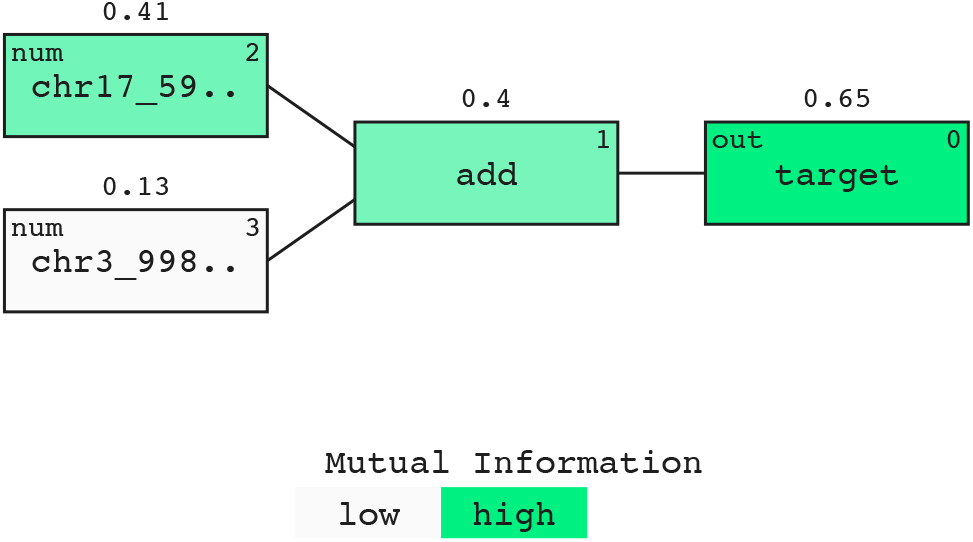
A representative model for predicting Hepatocellular Carcinoma. A prominent signal contribution from chr17_59473060_59483266 is found in all 10 models. The signal is expressed in terms of mutual information and displayed above the nodes in the model [35]

**Figure 8.**
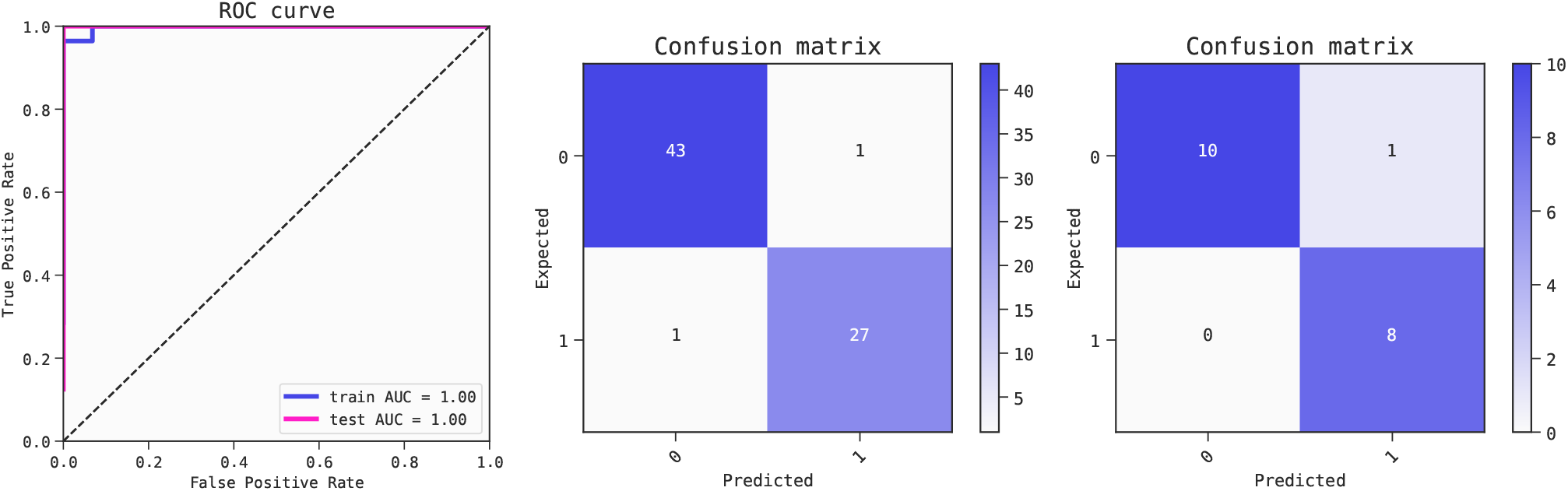
Metrics of the best model (ranked by BIC criterion) for predicting Hepatocellular Carcinoma. The model is robust as shown by the performance of the training set (AUC 1.0) compared to the test set (AUC 1.0). ROC curves (left) and confusion matrices for training set (center) and test set (right).

Using 5-fold cross-validation, the QLattice top models yielded an average performance of AUC = 0.93 (mean of the five folds, with a standard deviation of 0.01).

#### Model Interpretation

As can be seen in Fig. 9, the primary separator of the two features in the selected model is chr17_59473060_59483266. Individuals who do not have cancer have stable, low levels of methylated alleles, while individuals with cancer generally have higher, more variable levels of this trait. In addition, we find that some cancer individuals have low levels of chr17_59473060_59483266. Furthermore, from the 2d plot of partial dependence in Fig. 9 we can also see that low values of both chr17_59473060_59483266 and chr3_9987895_9989619 can be used to identify cancer individuals. This dynamic is captured well in the 2d partial dependence plot of Fig. 9. This is an easily understood model, two genes interacting, generating a top performing model.

**Figure 9.**
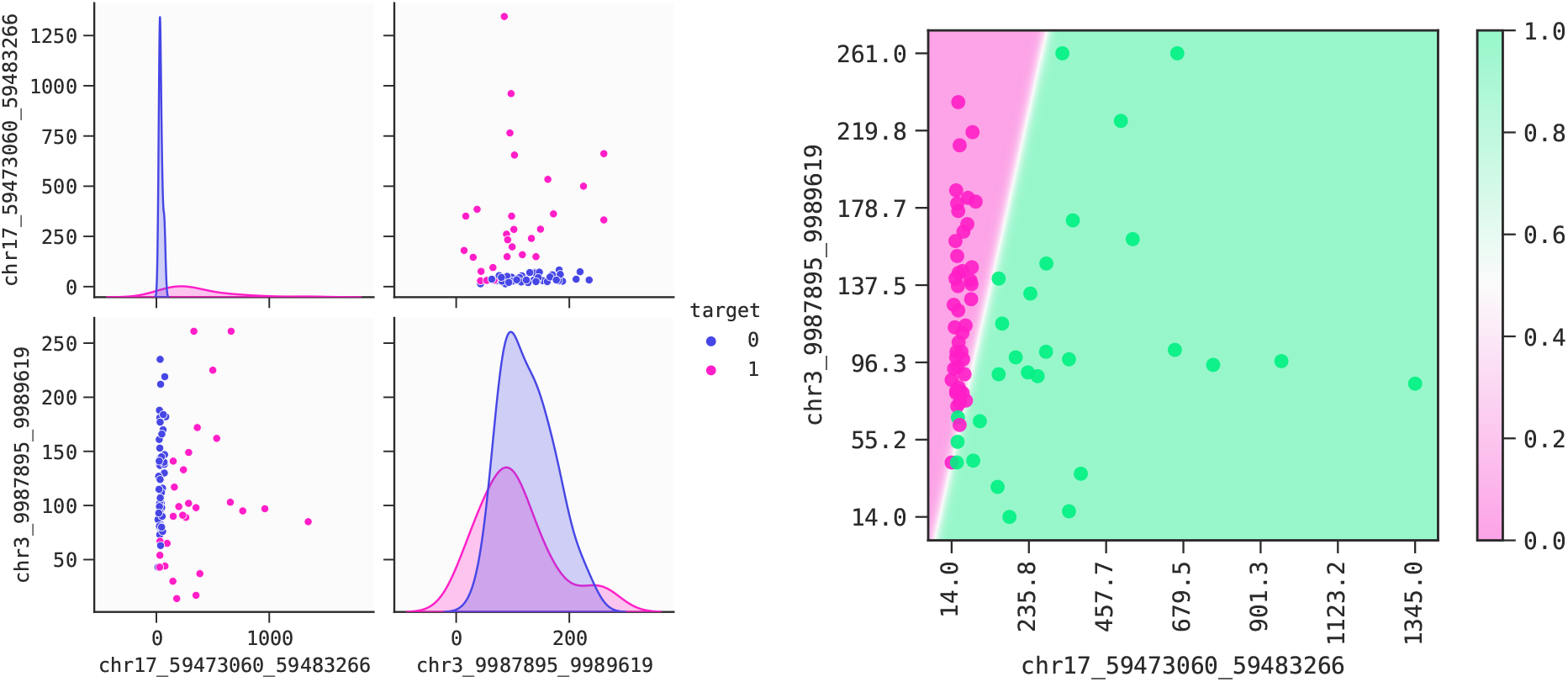
Left: HCC. Pairplot for features in the selected model. Right: 2d response of the model predictions, with train data overlaid. The decision boundary separates the two outcome areas.

**Figure 10.**
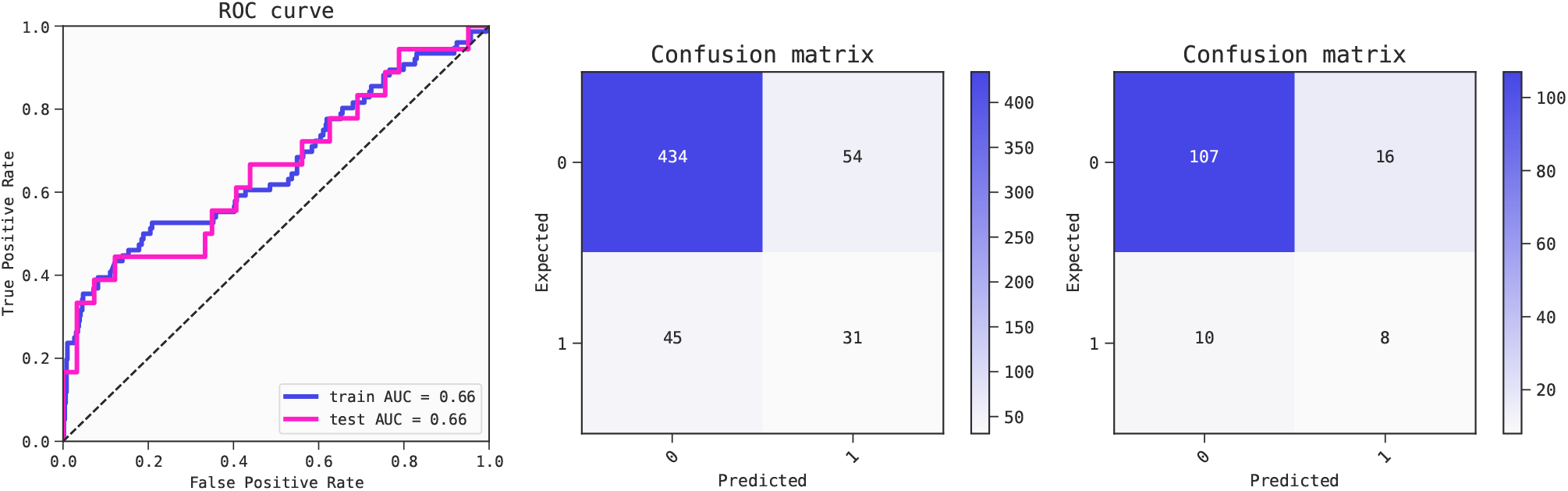
Metrics of the best model of the first fold (ranked by BIC criterion) for predicting Breast Cancer outcomes. The model is not overfitted as shown by the performance of the training set (AUC 0.66) compared to the test set (AUC 0.66). ROC curves (left) and confusion matrices for training set (center) and test set (right).

The models that were generated (Table 3) perform equally well. Aside from chr17_59473060_59483266 all models contain different secondary features and thus there could be molecular substitutes among the other features. To show whether there is multicollinearity, an overview of other correlated features is given in the correlation heatmap Fig 14. The figure shows whether the selected model feature belong to a group of highly correlated features. If this is the case, we can most likely replace this one feature with another from the same group and achieve similar model performance. In this case, the QLattice achieves high performance by selecting one feature from each main variance group in the dataset.

Instead of using dimensionality reduction like PCA or similar methods to group features with similar variance into single features, the QLattice selects representatives from each variance group. The representative that performs best in combination with the other features in the training dataset is selected.

### Multi-omics: Breast cancer

#### Background

Breast cancer is the most common cancer in women, worldwide. There are two main types of breast cancer, ductal and lobular carcinoma. The cancers can be classified as invasive or noninvasive. The noninvasive forms are often referred to as ductal carcinoma in situ (DCIS) and lobular carcinoma in situ (LCIS). Even though there are significantly different risks between patients, currently all lesions are treated. This can lead to excessive treatment of the condition in many patients. To complicate matters, breast cancer patients at similar stages of progression can have significantly different treatment responses and survival outcomes [46, 47].

In this case study, we explore a multi-omics dataset and identify potential regulatory interactions across omics-types (copy numbers (cn), somatic mutations (mu), gene expression (rs), protein expression (pp)) that could explain and predict survival outcomes of breast-cancer patients. We benchmark the QLattice models with a random forest and show that in addition to revealing interactions the QLattice performs as well as complex “black-box” models. The data set was obtained from Ciriello et al. [31] and contains multi-omics data from 705 breast tumor samples.

#### Two feature models analysis

Upon running the QLattice on different partitions of the data, one can expect different models being selected. These models bring similar albeit complementary insights, as they are able to see different sub-samples of the data. In this case, we obtained diverse models by keeping the lowest BIC-scoring ones from each partition of our cross-validation scheme.

To maximize interpretability, we started by exploring simple models that allowed for a maximum of two features. The mean test AUC for the best models of all folds was 0.635, with a standard deviation of 0.070. Equations (1) contain the best model (ranked by BIC) for each of the five folds.

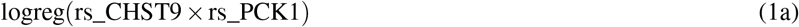

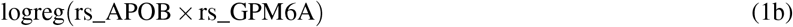

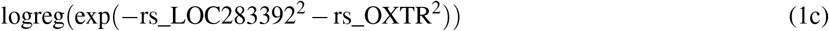

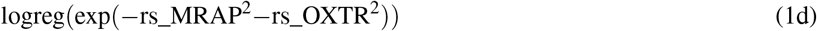

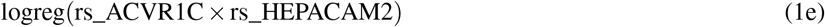

Two of the expressions correspond to a bivariate normal distribution, while the others have a multiplicative interaction as shown in equations (1). All the chosen features in the models above are measurements of gene expression.

From the Pearson correlation heatmap in Fig. 11, we observe that all five models contain a gene-expression feature from the group with highest pairwise Pearson correlation: *rs_PCK1, rs_MRAP, rs_LOC283392, rs_APOB* and *rs_ACVR1C*; their correlation values range from 0.774 to 0.835. Then these features are each combined in a non-linear interaction with the remaining gene expression variables.

**Figure 11.**
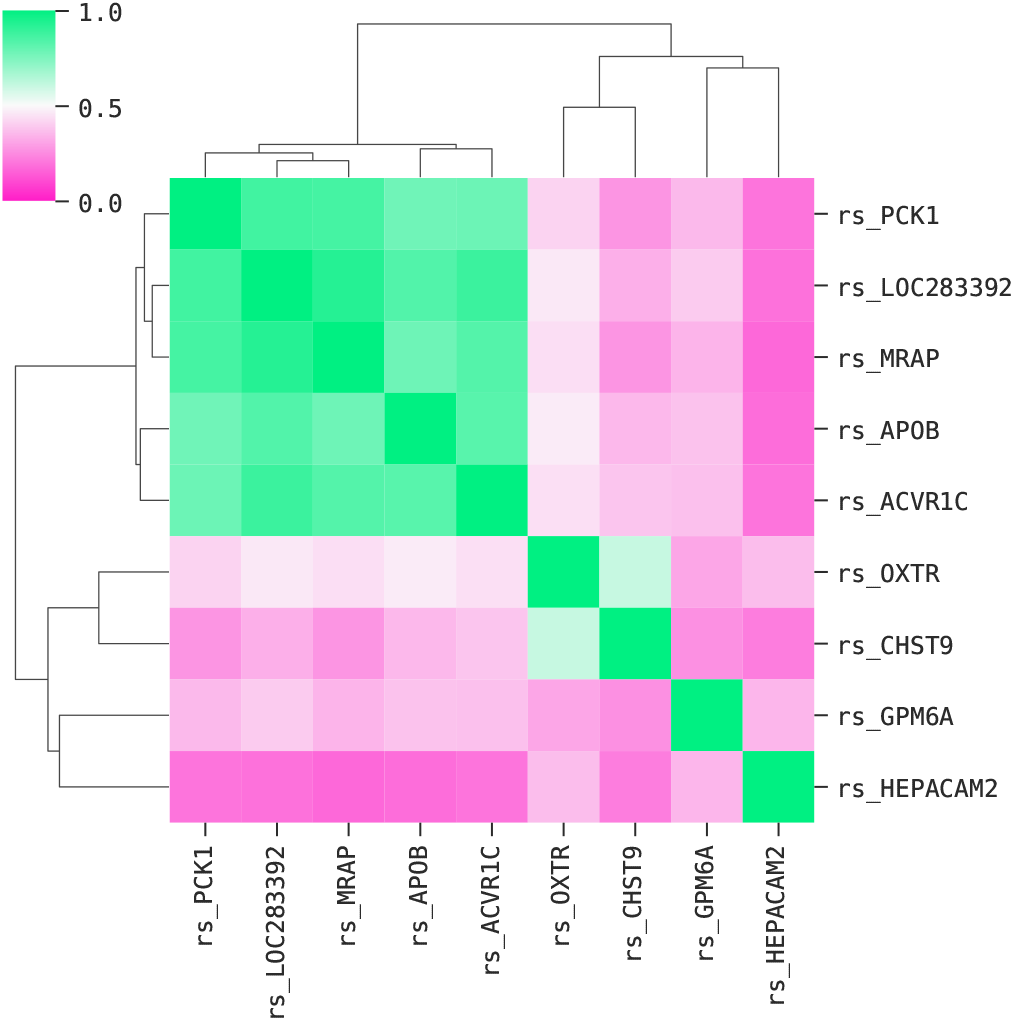
Pairwise Pearson correlation (absolute value) heatmap of the gene expression features in the models shown in equations (1).

Pairwise correlation gives a measure of the similarity between the input features. In addition, one can calculate the correlation between input features and the output variable, as shown in Table 4. The latter gives a measure of the relevancy of the input features relative to the output. Note in Table 4 that *rs_PCK1, rs_MRAP, rs_LOC283392, rs_APOB* and *rs_ACVR1C* are the features with highest relevance in this group. Therefore, akin to the HCC case, the models yielded by the QLattice combine a gene expression variable with high relevance with another gene expression with low similarity score.

**Table 4.** Pearson correlation between gene expression features and the output *vital*.*status*, the associated p-values, and the p-values adjusted for multiple hypothesis testing using the Bonferroni correction. Values computed using SciPy [48].

|  | Pearson corr. | p_value | p_value adj. |
| --- | --- | --- | --- |
| rs_APOB | 0.270 | 3.0e-13 | 5.9e-10 |
| rs_LOC283392 | 0.230 | 6.3e-10 | 1.2e-06 |
| rs_PCK1 | 0.225 | 1.6e-09 | 3.2e-06 |
| rs_MRAP | 0.214 | 1.0e-09 | 2.0e-05 |
| rs_ACVR1C | 0.206 | 3.1e-08 | 6.1e-05 |
| rs_OXTR | 0.194 | 2.0e-07 | 3.8e-04 |
| rs_CHST9 | 0.139 | 2.2e-04 | 4.3e-01 |
| rs_GPM6A | 0.116 | 2.0e-03 | 1.0e+00 |
| rs_HEPACAM2 | 0.051 | 1.8-01 | 1.0e+00 |

**Table 5.** Benchmarks for Alzheimer’s Disase dataset

|  | All features | MI top 10 | F top 10 | MI top 3 | F top 3 | LASSO Features |
| --- | --- | --- | --- | --- | --- | --- |
| QLattice | 0.929 | - | - | - | - | - |
| LASSO | 0.875 | - | - | - | - | - |
| Elasticnet | 0.657 | - | - | - | - | - |
| Random Forest | 0.943 | 0.953 | 0.955 | 0.934 | 0.951 | 0.982 |
| Gradient Boosting | 0.989 | 0.966 | 0.95 | 0.942 | 0.937 | 0.973 |

**Table 6.** Benchmarks for Insuline Response dataset

|  | All features | MI top 10 | F top 10 | MI top 3 | F top 3 | LASSO Features |
| --- | --- | --- | --- | --- | --- | --- |
| QLattice | 0.957 | - | - | - | - | - |
| LASSO | 0.963 | - | - | - | - | - |
| Elasticnet | 0.841 | - | - | - | - | - |
| Random Forest | 0.957 | 0.966 | 0.967 | 0.954 | 0.957 | 0.98 |
| Gradient Boosting | 0.971 | 0.969 | 0.969 | 0.945 | 0.95 | 0.975 |

**Table 7.** Benchmarks for Hepatocellular Carcinoma dataset

|  | All features | MI top 10 | F top 10 | MI top 3 | F top 3 | LASSO Features |
| --- | --- | --- | --- | --- | --- | --- |
| QLattice | 0.959 | - | - | - | - | - |
| LASSO | 0.959 | - | - | - | - | - |
| Elasticnet | 0.944 | - | - | - | - | - |
| Random Forest | 0.968 | 0.948 | 0.961 | 0.941 | 0.951 | 0.964 |
| Gradient Boosting | 0.975 | 0.959 | 0.95 | 0.946 | 0.916 | 0.963 |

**Table 8.** Benchmarks for Breast Cancer dataset

|  | All features | MI top 10 | F top 10 | MI top 5 | F top 5 | LASSO Features |
| --- | --- | --- | --- | --- | --- | --- |
| QLattice | 0.633 | - | - | - | - | - |
| LASSO | 0.568 | - | - | - | - | - |
| Elasticnet | 0.611 | - | - | - | - | - |
| Random Forest | 0.604 | 0.628 | 0.558 | 0.594 | 0.541 | 0.604 |
| Gradient Boosting | 0.570 | 0.621 | 0.541 | 0.591 | 0.59 | 0.594 |

Let us take a look at some of the models’ decisions boundaries depicted in Fig. 12. The bivariate gaussian function and the product between the gene expression features identify a “hotspot”, i.e., there is a particular range for these gene expressions that indicate whether a breast-cancer patient died or survived. Strikingly, these patients were predominantly suffering from ductal breast cancers. There seems to be a putative molecular interaction that is an important biomarker for ductal breast-cancer survival.

**Figure 12.**
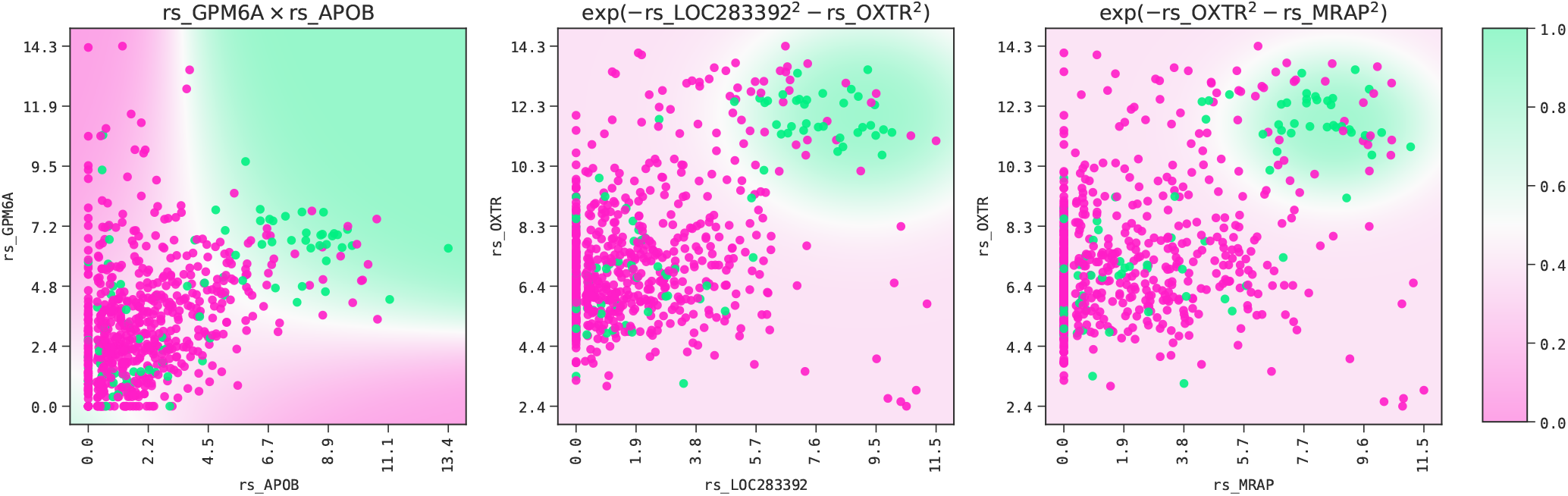
Decision boundary for three of the models at the head of each k-fold. Green indicates a higher probability of a death outcome, while pink represents the opposite.

**Figure 13.**
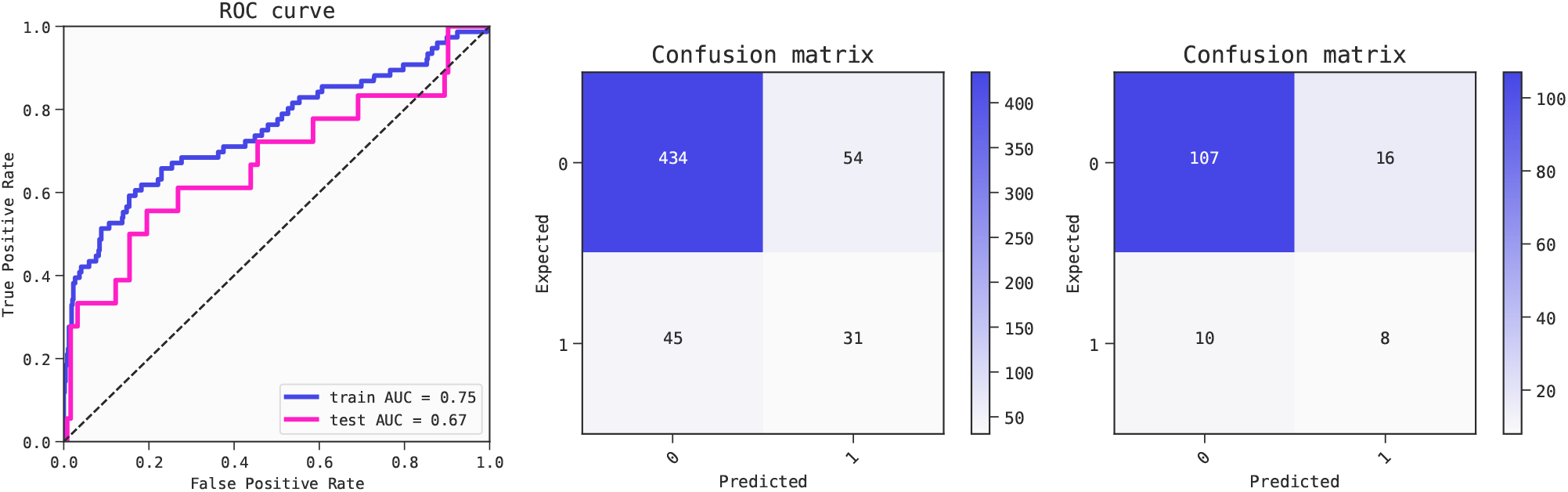
Metrics of the best model of the first fold (ranked by BIC criterion) for predicting Breast Cancer outcomes. The model shows some degree of overfitting as shown by the performance of the training set (AUC 0.75) compared to the test set (AUC 0.67). ROC curves (left) and confusion matrices for training set (center) and test set (right).

**Figure 14.**
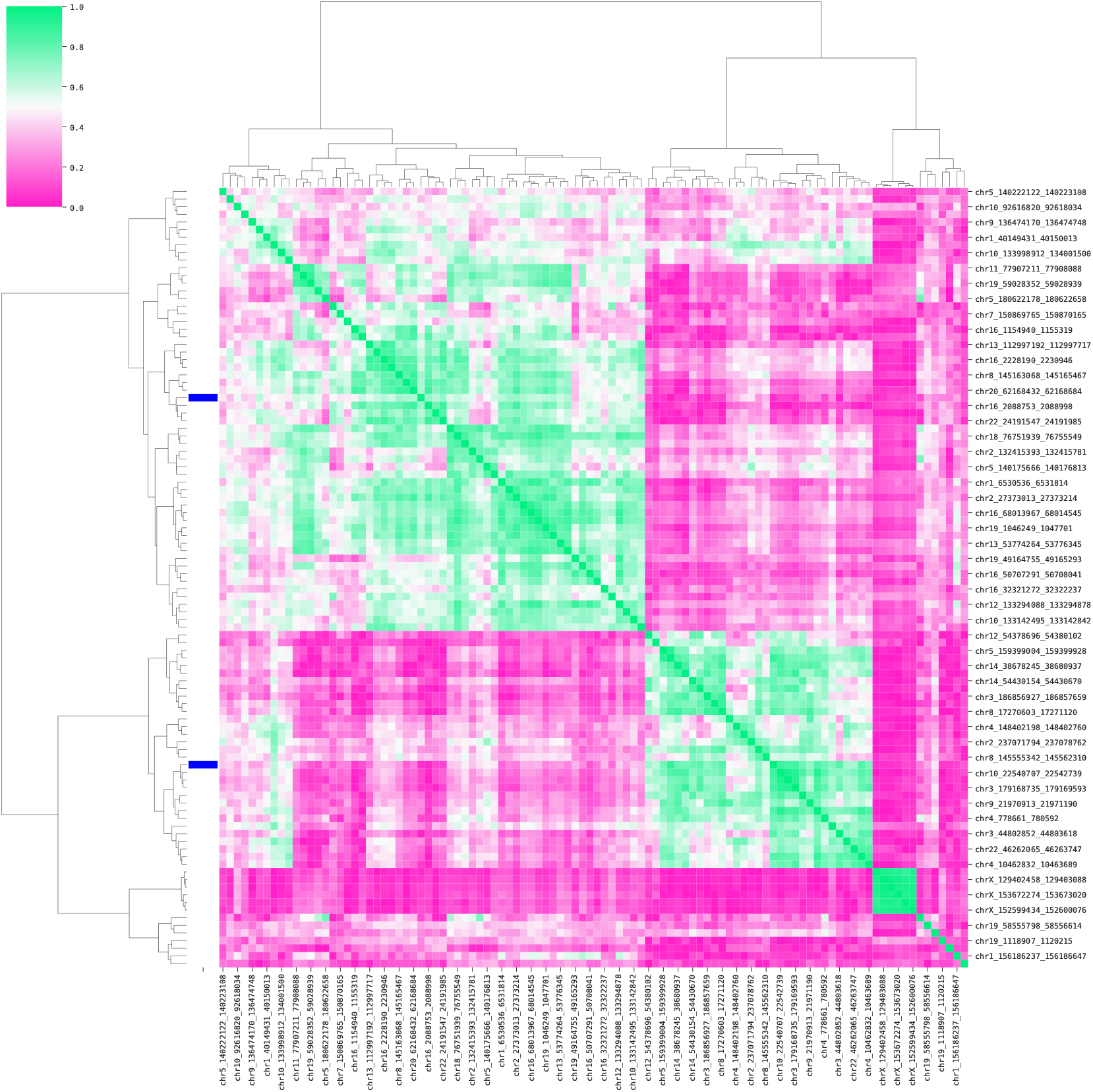
HCC correlation heatmap for pairwise correlations (in absolute value) between a random subset of 100 features including the two model features (blue bars on the left). The heavy green coloring confirms extensive multi-collinearity. The plot clusters features based on similarity measured by the Pearson correlation coefficient followed by sorting. The two main groups of linear feature variance are displayed by the top branches in the dendrogram. Clustermap function from Seaborn [27].

Lastly, the genes in the models depicted in equations (1) were found to have no relation to the age of the patients. The

Pearson correlation coefficient between gene expression and age was computed and the highest absolute Pearson correlation coefficient value between gene expression and age was 0.153 (p-value of 4 × 10^−5^).

#### Comparison with multi-omics models

Allowing models with higher complexity – more features and operations – can potentially unlock better performing models that mix different *omics*. To this end, we ran the same 5-fold cross-validation scheme allowing a maximum of five features. This should allow for any signal beyond the gene expression “hotspot” to be captured by the QLattice. The resulting average test AUC score on the best models is 0.671 with standard deviation of 0.040. This average result is certainly larger than test AUC of the two feature models, although both scores could be considered statistically compatible.

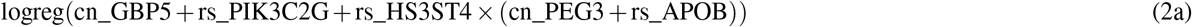

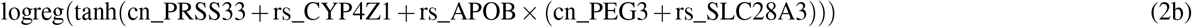

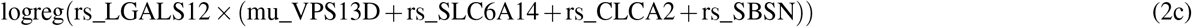

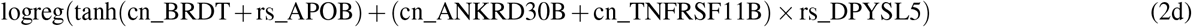

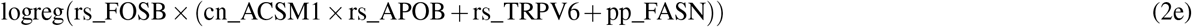

The average train AUC of the models on Eqs. 2 is 0.743 (*σ* = 0.020), which is significantly higher than the average train AUC of the two-features models, 0.683 (*σ* = 0.043). Since their mean test AUC scores are on par, the discrepancy in the training set implies that the more complex models depicted above tend to overfit when compared to the simpler gene expression models presented before. When it comes to the functional form of the models on Eqs. 2, it is interesting to note that they all possess a non-linear interaction between gene expression features (prefix *rs*). For most, this interaction is multiplicative, while for the model from Fold 3, the non-linear boundary is set by the tanh function of *rs_APOB*.

#### Comparison with random forest

Lastly, to get a better sense of the performance of the QLattice, we compare it with a widely used “black-box” model: the random forest. We use the implementation by `scikit-learn`, and tune its hyperparameters and estimate its performance using a “nested” cross-validation scheme [49, 50]. The best parameters lay around `n_estimators` = 50 and `max_depth` = 4 for the different folds, and the average performance is an AUC of 0.604 with standard deviation of 0.106, on par with the QLattice. This is a very remarkable result, considering that the QLattice is only using two features while the random forest uses potentially all of them. A benchmark with other algorithms and feature selection techniques can be found in the supplementary material.

Taking into consideration how the multi-omics models in Eqs 2 tend to overfit and the random forest result in comparison to the QLattice, we can conclude that the models in Eq. (1) reveal the core patterns in the data. In summary, the interaction of two gene expression variables allows for the identification of a “hotspot” where the probability of a poor outcome of the disease is high. One of the genes in the model belongs to a group of genes with pairwise Pearson correlation above 0.7, while the other is taken from the remaining pool of variables. A possible next step in the study of this data is to pinpoint the combinations of gene expression variables that best predict *vital*.*status*.

## Conclusions

Given the large amounts of data being generated and a need for more efficient treatment regimens, predictive analytics in the clinic is gaining traction. A range of methods exist that can predict a certain outcome based on omics data, however there is a scarcity of interpretable alternatives. Here, we showed that we can identify simple yet highly predictive and explainable biomarker signatures by combining sophisticated feature selection with a powerful model search algorithm. Due to the small number of features, the models are robust and can be readily interpreted. This makes them a valuable starting point for researchers and clinicians who are looking to find new and biomarker signatures while learning about the underlying interactions in the data.

## Supplementary material

### Code and data

All code and data used to generate the models and plots discussed in this work can be found in https://github.com/abzu-ai/QLattice-clinical-omics.

### Benchmarks

In order to have a fair and comprehensive benchmark for the QLattice, we trained and evaluated other machine learning algorithms in combination with different feature selection techniques on all four datasets. We used the nested cross-validation scheme detailed in [49, 50] for hyperparameter tuning and performance estimation. The algorithms tested are LASSO, Elastic Net, Random Forest and Gradient Boosting, and the feature selection techniques are LASSO, top N features based on Mutual Information, and top N features based on the F-test (where N is either ten or the number of features used by the QLattice models). All implementations are from scikit-learn [51]. Performance benchmarks on regression problems for the QLattice and other algorithms can be found in a previous work [14].

Upon running the Friedman test (implementation by SciPy [48]) on the best performing models (QLattice, and Random Forest and Gradient Boosting in combination with feature selection based on Mutual Information and LASSO), we obtain a p-value of 0.62. Hence, we reject the hypothesis of one model being consistently better than the rest.

## Acknowledgements

We would like to acknowledge Jonas Elsborg and Caroline Linnea Elin Lennartsson for their contributions to the manuscript.

## Author contributions statement

V.S.H, M.M., M.T.I., M.S., S.D. and N.J.C. analysed the data and wrote the manuscript. V.S.H. and M.T.I. performed the cross validation of the results. All authors reviewed the manuscript.

